# EyeLoop: An open-source, high-speed eye-tracker designed for dynamic experiments

**DOI:** 10.1101/2020.07.03.186387

**Authors:** Simon Arvin, Rune Rasmussen, Keisuke Yonehara

## Abstract

Eye-tracking is a method for tracking the position of the eye and size of the pupil, often employed in neuroscience laboratories and clinics. Eye-trackers are widely used, from studying brain dynamics to investigating neuropathology and disease models. Despite this broad utility, eye-trackers are expensive, hardware-intensive, and proprietary, which have limited this approach to high-resource facilities. Besides, experiments have largely been confined to static open-loop designs and post hoc analysis due to the inflexibility of current systems. Here, we developed an open-source eye-tracking system, named EyeLoop, tailored to dynamic experiments. This Python-based software easily integrates custom functions via a modular logic, tracks a multitude of eyes, including rodent, human, and non-human primate eyes, and it operates well on inexpensive consumer-grade hardware. One of the most appealing applications of EyeLoop is closed-loop experiments, in which the eyes evoke stimulus feedback, such as rapid neuronal optogenetic stimulation. By using EyeLoop, we demonstrate its utility in an open-loop, a closed-loop, and a biomedical experiment. With a remarkably low minimal hardware cost amounting to 29 USD, EyeLoop makes dynamic eye-tracking accessible to low-resource facilities, such as high schools, small laboratories, and small clinics.

## Introduction

At every moment, the brain uses the senses to produce increasingly complex features along its computational hierarchy [1, 2]. Our every-day behaviors, such as navigating in traffic, are directed in large part by our sensory input, that is, what we see, hear, feel, etc. [3]. The senses are incredibly dynamic instruments that reflexively facilitate their own reception, e.g., by head movements directed by attention [4], whiskers scanning the immediate proximity [5], the nose sniffing aromas [6], or by deformation of the ears [7]. The eyes, in particular, engage in sensory facilitation: For example, the optomotor reflex detects perturbations of visual flow to avoid collisions in insects [8] and elicits stabilizing head movements in mice [9]. Recently, combined eye-head movements in free-roaming mice were shown to re-align the visual axis to the ground plane [10] (see also [11]).

Eye-trackers are widely used, from studying brain dynamics to investigating neuropathology and disease models. Despite this broad utility, eye-trackers are expensive, hardware-intensive, and proprietary, which have constrained its use to high-resource facilities. In addition, due to the inflexibility of current systems, experiments have largely been confined to static open-loop designs and post hoc analysis, failing to capture the dynamicity of the systems they seek to elucidate.

Here, we developed an open-source eye-tracking system – EyeLoop – tailored toward investigating sensory dynamics. EyeLoop enables low-resource facilities access to eye-tracking and encourages the integration of custom functions through a modular logic.

## Design and implementation

### Software overview

EyeLoop is based on the versatile programming language, Python 3. Contrary to other frameworks used for eye-tracking, e.g., LabView [12] and MATLAB [13], or proprietary systems, e.g., ISCAN [14, 15], Python is open-source and has recently seen a surge in popularity, particularly owing to its outstanding modularity and standard library [16]. In the same spirit, EyeLoop experiments are built simply by combining modules (Fig 1).

**Fig 1.**
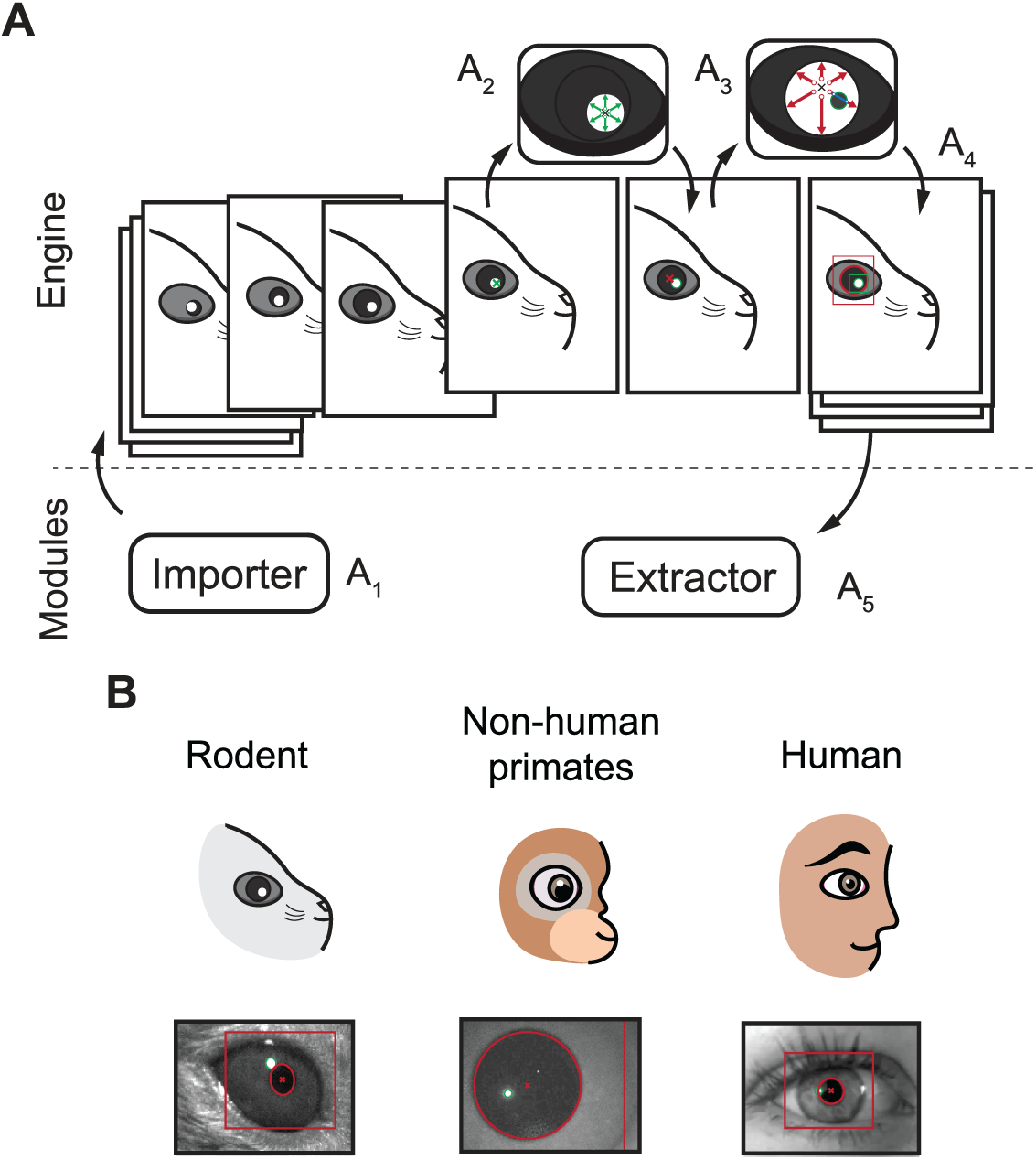
Schematic overview of the EyeLoop algorithm and its applications. (A) Software overview. The engine exchanges data with the modules. The importer module imports camera frames in a compatible format (A_1_). The frame is binarized and the corneal reflections are detected by a walk-out algorithm (A_2_). Using the corneal reflections, any pupillary overlap is removed, and the pupil is detected via walk-out (A_3_). Finally, the data is formatted in JSON and passed to all modules, such as for rendering (A_4_), or data acquisition and experiments (A_5_). (B) EyeLoop accepts a variety of animal eyes, including rodents, non-human primates, and humans, by employing distinct mathematical models for each type of eye.

EyeLoop consists of two domains: the engine and the modules. The engine detects the pupil and corneal reflections, whereas the modules import or extract data from the engine. *Extractor* modules are used in data acquisition or to produce experimental schemes, and *importer* modules are used to import video sequences. Notably, the graphical user interface is a module, too. This high modularity improves compatibility across software versions and camera types, and eases troubleshooting. Due to the modular user interface, EyeLoop is readily scaled to any hardware capacity: Minimal interfaces may be used in low resource facilities, such as classrooms, and comprehensive interfaces may be used in laboratories combined with advanced techniques, such as imaging, electrophysiology, or optogenetics.

The engine processes each frame of the video sequentially (Fig 1A_1_). First, the user selects the corneal reflections, then the pupil (Fig 1A_2-4_). The frame is binarized, filtered, and smoothed by a gaussian kernel (Fig 1A_2,3_). Then, the engine utilizes a walk-out algorithm to detect contours [12]. This produces a matrix of points, which is filtered to discard bad matches. Finally, the shape is parameterized by a fitting model: either an ellipsoid (suitable for rodents, cats, etc.) [17-19], or circle model (human, non-human primates, rodents, etc.) [20]. The target species is readily changed (“—model”).

Users build experiments by connecting extractors to EyeLoop. Extractors contain an instantiating constructor called at startup, and a fetch function called every time-step that has access to all eye-tracking data in real-time (Fig S1).

**Fig S1.**
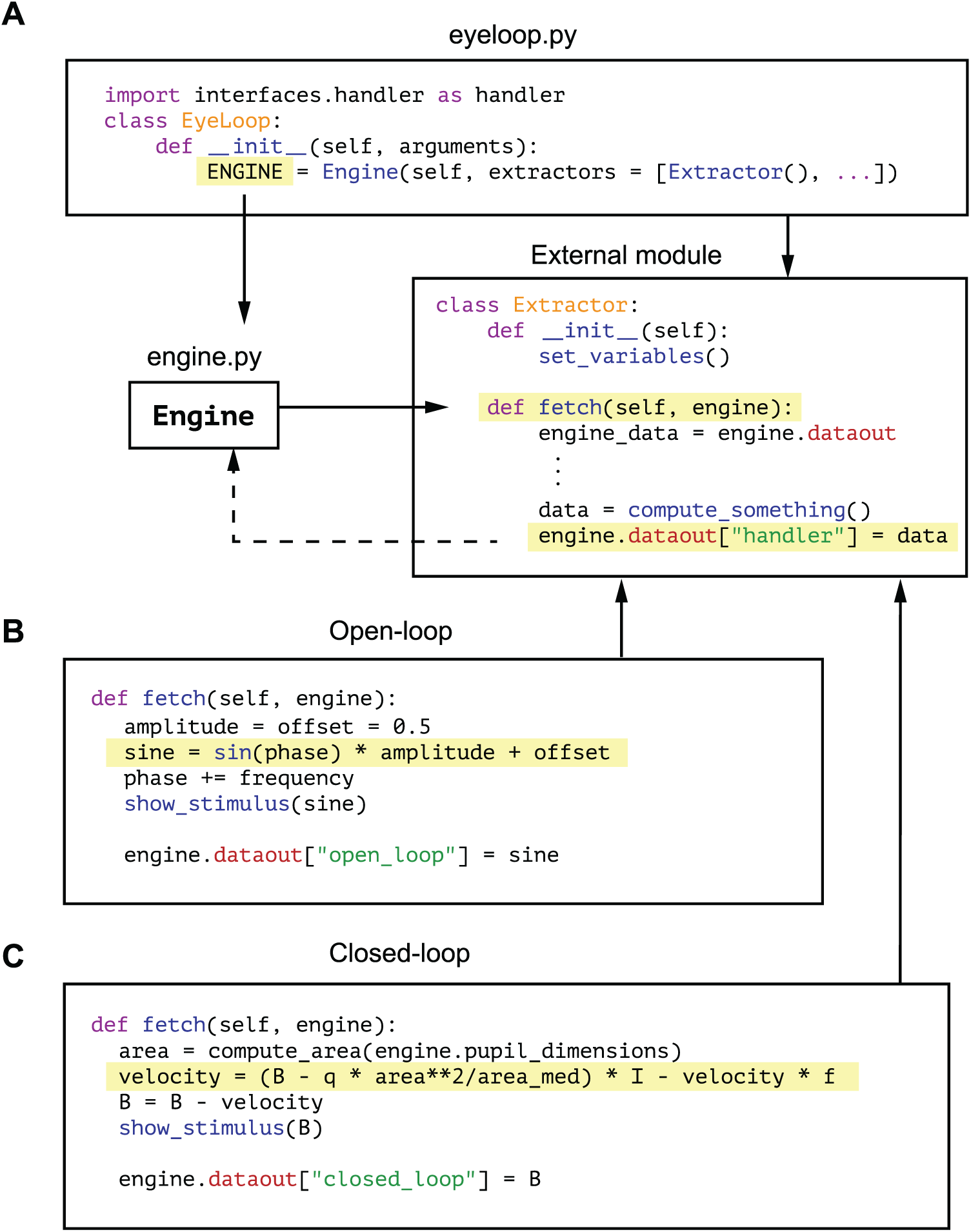
Schematic overview of EyeLoop’s programmatic logic. (A) Using EyeLoop’s main file, eyeloop.py, users can add custom modules. (B) Simplified script for the open-loop experiment; the sine function dictates the monitor brightness. (C) Simplified script for the closed-loop experiment; the monitor brightness depends on the pupil area, and the pupil area depends on the monitor’s brightness.

### Open-loop setting

To demonstrate the utility of EyeLoop in open-loop experiments, we designed an extractor class that modulates monitor brightness, *B*, based on the phase of the sine function (Fig S1B):

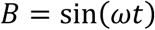

Where ω is the angular frequency, and *t* is the frame number. Using this design, we sought to confirm that the pupillary constriction speed dominates the speed of dilation in mice, mirroring findings in humans (Fig 2) [21].

**Fig 2.**
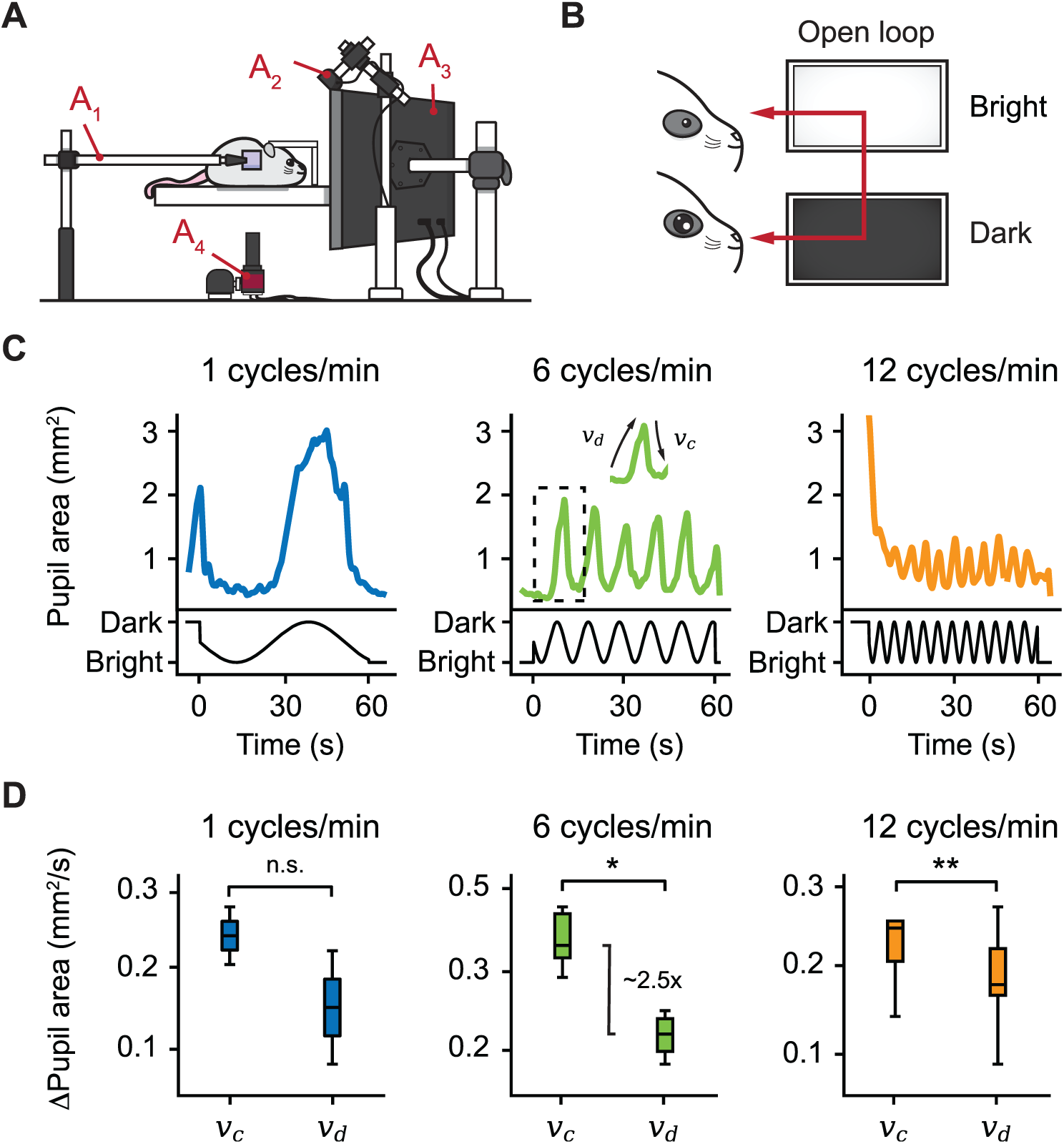
Open-loop experiments confirm constriction speed dominance in mice. (A) The setting used for eye-tracking in mice. A hot mirror is positioned beside the mouse and above the camera (A_1_,_4_). A monitor displaying the visual stimulus is positioned facing the mouse (A_3_), while a near-infrared light source is placed in the back (A_2_). (B) Open-loop experiment. A sine function is mapped onto the brightness of a monitor, producing oscillations in the pupil area. (C) Example plots from three open-loop experiments with frequencies 1, 6, and 12 cycles/min. (D) Constriction speed (v_c_) and dilation speed (v_d_) for each frequency, calculated using the first derivative of the pupil area plots (n = 3 mice). Centerline is median, box limits are 25th and 75th percentiles, and whiskers show the minimum and maximum values. \**P* < 0.05, \*\**P* < 0.01, n.s., not significant, Wilcoxon signed-rank test.

Pupil size is modulated by a special class of intrinsically photosensitive cells in the retina that project to the upper midbrain and modulate pupil size in the pupillary light reflex [22, 23]. To test our hypothesis, we ran an array of three frequencies ranging from 1 to 12 cycles/min. As the phase of the sine function shifts, monitor brightness changes correspondingly. First, as expected, our findings suggest that pupil size entrains to monitor brightness by inverse proportionality, a consequence of the pupillary light reflex (Fig 2C). Second, using the pupil area’s first derivative, we confirm that constriction speed is greater than dilation across all trials [21]. Notably, while both high and low-frequency trials exhibit modest constriction speeds, approximately 1.5 times greater than dilation, we observed a significant 2.4-fold greater constriction speed for the mid-frequency trials. Thus, our findings lend support to mathematical models predicting that the light response depends on the statistics of the light stimulus itself [24].

### Closed-loop setting

One of EyeLoop’s most appealing applications is closed-loop experiments (Fig 3). To demonstrate this, we designed an extractor class to use the pupil area to modulate the brightness of a monitor, in effect a reciprocal feedback loop (Fig S1C): Here, the light reflex causes the pupil to dilate in dim settings, which causes the extractor to increase monitor brightness. In turn, this causes the pupil to constrict, causing the extractor to decrease brightness and return the experiment to its initial state.

**Fig 3.**
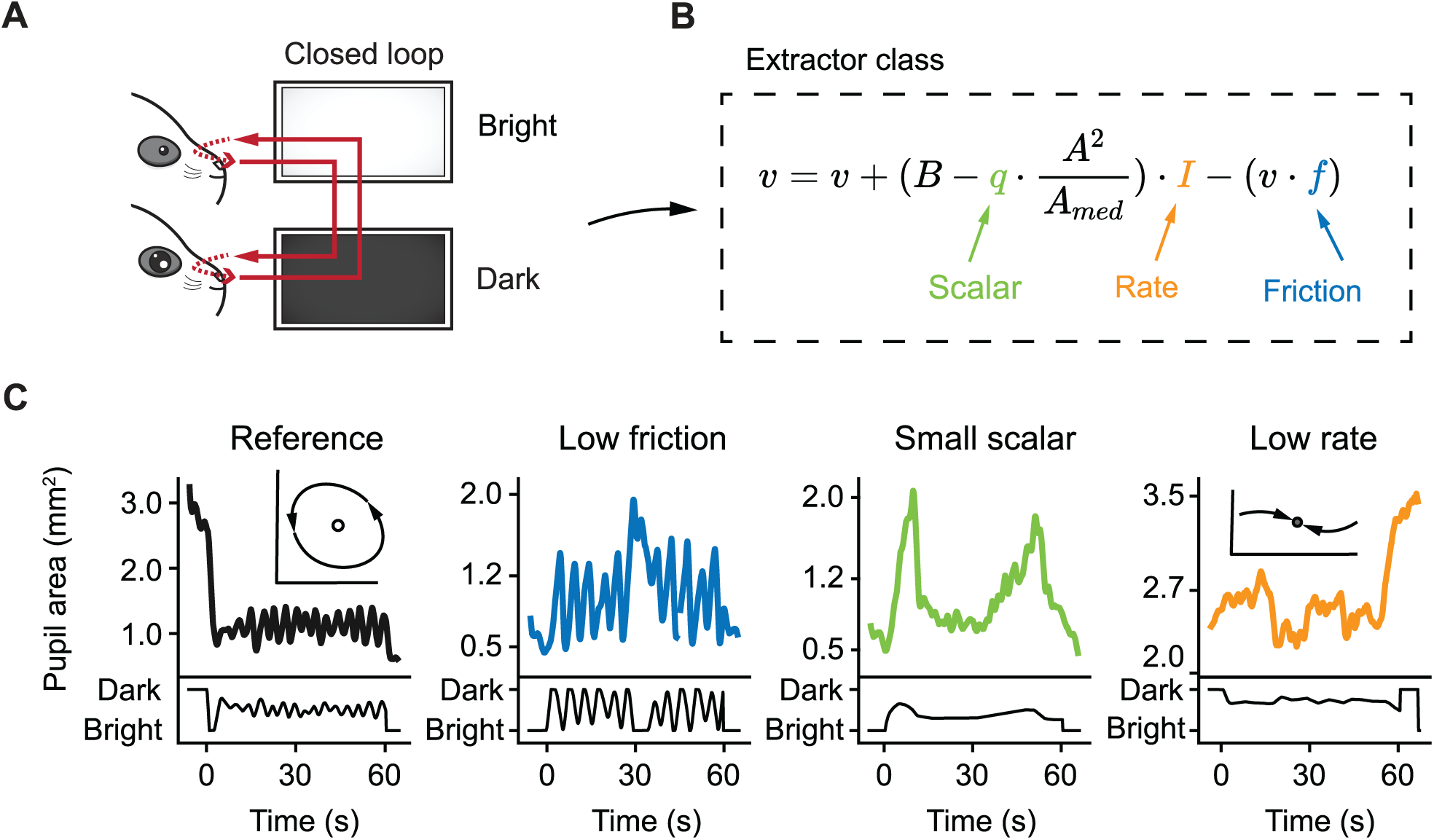
Closed-loop experiment exhibiting properties of dynamical systems. (A) Closed-loop experiment using reciprocal feedback. Monitor brightness is controlled as a function of the pupil area. (B) State velocity, *v*, depends on pupil area, A. (C) Four trials of the closed-loop experiment with differing parameters showing distinct dynamic behavior.

The brightness formula contains four critical variables (Fig 3B): The rate of change, *I*, which is dependent on the pupil area, and its scalar, *q*. The velocity, *v*, which applies the rate of change to monitor brightness, and, the velocity friction, *f*, which decays the velocity towards zero. Interestingly, by varying these parameters, we observe behaviors characteristic of dynamical systems: For the reference and the slow decay trials, we find emergent limit-cycle oscillations (Fig 3C). This dynamic is dramatically impaired by a small scalar, and abolished in the low rate trials. These findings illustrate how a simple closed-loop extractor generates self-sustaining dynamics emerging from the eyes engaging with the system, and the system engaging with the eyes.

### Optokinetic reflex in congenital nystagmus model versus wild-type mice

Often, neuropathologies, such as latent brain hemorrhage and Horner syndrome, cause eye abnormalities. Similarly, patients suffering from congenital nystagmus lack the optokinetic reflex, a type of eye movement for gaze stabilization evoked by rotation. To demonstrate how EyeLoop may be applied as a diagnostic tool, we here confirm previous findings on *Frmd7* hypomorphic mice by showing that *Frmd7* knockout mice lack the horizontal optokinetic reflex; similar to *Frmd7*-mutated congenital nystagmus patients [25]. To this end, we simulated rotation using a bilateral drifting grating stimulus (Fig 4A). As expected, for wild-type mice, the optokinetic reflex was faithfully evoked (Fig 4B). Conversely, this reflex was absent in *Frmd7* knockout mice, verifying the disease phenotype (Fig 4C).

**Fig 4.**
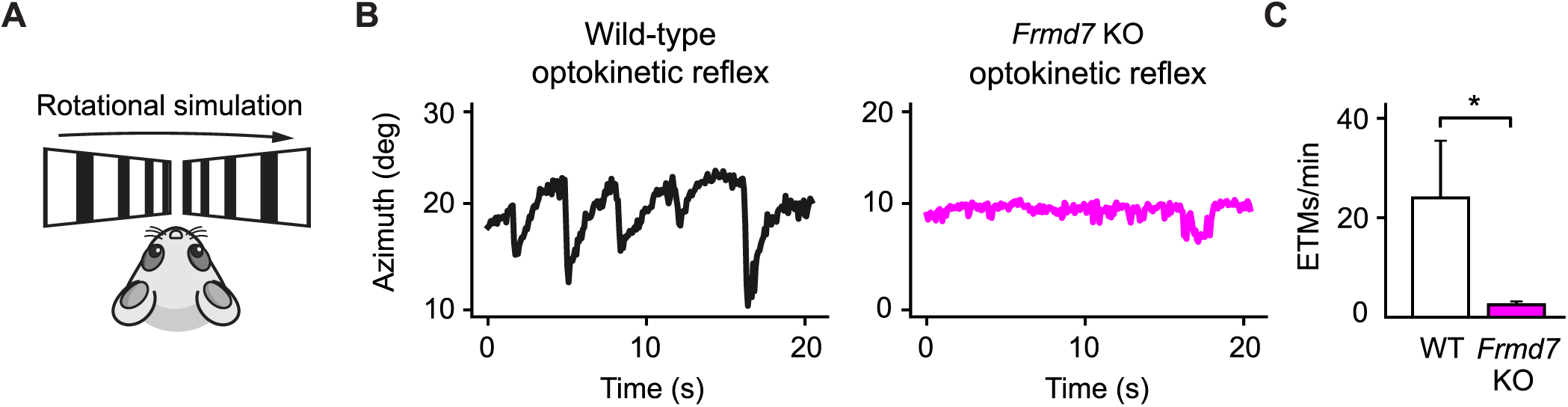
The horizontal optokinetic reflex is absent in *Frmd7* knockout mice. (A) Rotational motion simulation using gratings drifting in parallel along the horizontal axis. (B) Eye movements evoked by the optokinetic reflex in wild-type and *Frmd7* KO (knockout) mice in response to drifting grating stimulation. The azimuth represents the horizontal angular coordinate of the eye. (C) The optokinetic reflex was quantified as eye-tracking movements per minute (ETMs), computed by thresholding the first derivative of eye movements. WT; n = 3, *Frmd7*; n = 3. Error bars show standard deviation. \**P* < 0.05, Wilcoxon signed-rank test.

### Diverse applications

EyeLoop fills an important gap as a tool to investigate the role of the eyes in brain dynamics (Fig 5). Sensory integration is complex, and often the eyes play an instrumental role in its orchestration. Eye-tracking during sensory exploration carries enormous information on how senses are used by the brain: For example, during fast whole-body rotation, the eyes act to stabilize the gaze via the vestibulo-ocular reflex integrating both vestibular and visual signals [26]. Despite the known complexities of sensory computations, visual experiments are usually aimed at strictly monitoring the eyes [27], or at applying one-sided perturbations, such as an open-loop stimulus [14]. EyeLoop enables researchers to use the eyes as active elements in their experiments, for example to investigate neuroplasticity by timing learning cues to distinct arousal states based on pupil size [28-31]. Future experiments could use eye movements to silence or stimulate specific neuronal populations via optogenetics to investigate the causal relationship between neuronal activity and the endogenous parameters by which the neural system operates [32].

**Fig 5.**
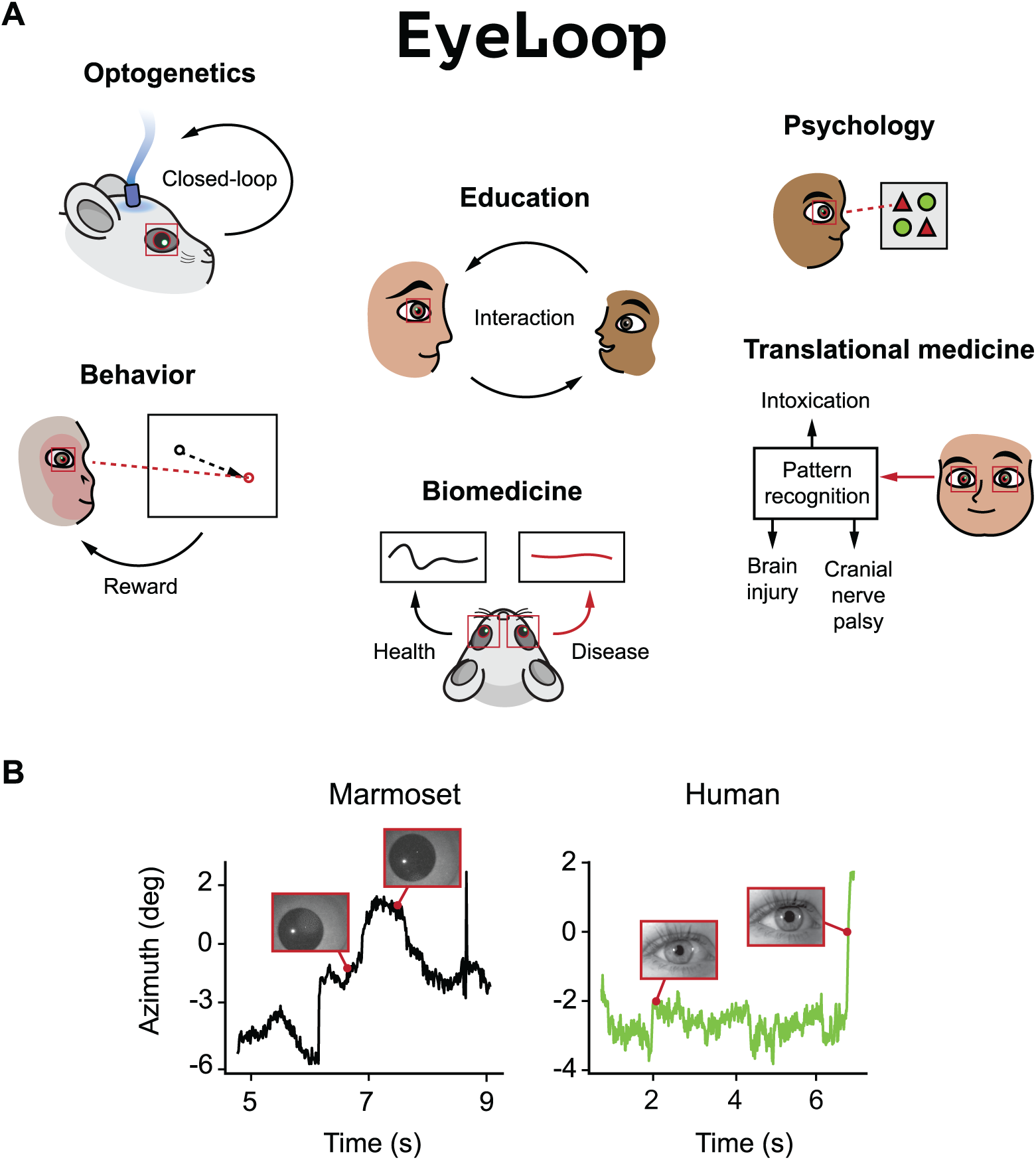
EyeLoop can be used in a multitude of applications. (A) Example applications where EyeLoop may be used in the future. (B) Eye-tracking traces for common marmoset and human sporadic eye movements.

In clinical medicine, the behavior of the eyes is increasingly becoming a matter of interest. Indeed, extensive studies on eye movements in healthy and ill patients have revealed that disorders of the brain often produce distinct abnormalities of the eyes [33, 34]. Accordingly, EyeLoop offers clinical researchers a system for tracking eye movements in human and non-human primates (Fig 5B) [35]. Due to its flexibility, custom functions are easily integrated and could be applied in the neurological setting to improve diagnostic sensitivity and patient care, e.g., using automatically recognized eye abnormalities. Similarly, assessment of pupil size could aid the diagnosis of neurological disorders, such as Horner syndrome and oculomotor palsy [36, 37], or reveal intoxications, e.g., pin-point pupils characteristic of an opioid overdose [38]. In biomedicine, disease model animals, such as the congenital nystagmus mouse, could be investigated and characterized using EyeLoop. This provides an easy and non-invasive method for identifying novel disease models, and for verifying the phenotype of existing models.

With a remarkably low minimal hardware cost amounting to 29 USD (Yiduore 480P camera: 4.79 USD; Univivi Infrared Illuminator: 24.99 USD), EyeLoop enables any small capacity facility, such as small laboratories or high-school classrooms, to apply eye-tracking in research and education. In the spirit of open-source collaboration, users are encouraged to design new modules and contribute to the ongoing development of EyeLoop.

## Acknowledgments

We thank Jude Mitchell for video footage of marmoset pupil, Zoltan Raics for developing our visual stimulation system, and Bjarke Thomsen and Misugi Yonehara for technical assistance. R.R. was supported by Lundbeck Foundation PhD Scholarship (R230-2016-2326). K.Y. was supported by Lundbeck Foundation (DANDRITE-R248-2016-2518; R252-2017-1060), Novo Nordisk Foundation (NNF15OC0017252), Carlsberg Foundation (CF17-0085), and European Research Council Starting (638730) grants.

## Author contributions

S.A. and K.Y. conceived and designed the project. S.A. developed the software. S.A. and R.R. performed the experiments. S.A. analyzed the data. S.A. and K.Y. interpreted the data and wrote the paper.

## Materials and correspondence

Correspondence and requests for materials should be addressed to corresponding author, Keisuke Yonehara.

## Methods

### Ethics statement

All experiments on mice were performed according to standard ethical guidelines and were approved by the Danish National Animal Experiment Committee (2020-15-0201-00452). No experiment on non-human primate was conducted in this study; the marmoset video was kindly provided by Jude Mitchell (University of Rochester).

### Availability

All source code files are freely available online under the GNU General Public License v3.0 at the Yonehara Lab website (http://www.yoneharalab.com) and the Git repository (git: https://github.com/simonarvin/eyeloop). Here, we provide instructions on the installation and utilization of the system and full examples for testing. The minimum software requirements are Python 3, the machine vision module, OpenCV, and the numerical computing module, NumPy.

## Supplementary materials

### Experimental animals

Wild-type control mice (C57BL/6J) were obtained from Janvier Labs. *Frmd7* knockout mice are homozygous female or hemizygous male *Frmd7*^*tm1b(KOMP)Wtsi*^ mice, which were obtained as *Frmd7*^*tm1a(KOMP)Wtsi*^ from the Knockout Mouse Project (KOMP) Repository, Exon 4 and neo cassette flanked by loxP sequences were removed by crossing with female Cre-deleter *Edil3*^*Tg(Sox2-cre)1Amc*/J^ mice (Jackson laboratory stock 4783) as confirmed by PCR of genome DNA and maintained in a C57BL/6J background. Experiments were performed on 3 male and female wild-type control mice, and 3 male and female *Frmd7*^*tm*^ mice. All mice were between two and four months old. Mice were group-housed and maintained in a 12-hour/12-hour light/dark cycle with *ad libitum* access to food and water.

### Head-plate implantation

Surgeries and preparation of animals for experiments were performed as previously described [39]. Mice were anesthetized with an intraperitoneal injection of fentanyl (0.05 mg/kg body weight; Hameln), midazolam (5.0 mg/kg body weight; Hameln) and medetomidine (0.5 mg/kg body weight; Domitor, Orion) mixture dissolved in saline. The depth of anesthesia was monitored by the pinch withdrawal reflex throughout the surgery. Core body temperature was monitored using a rectal probe and temperature maintained at 37–38°C by a feedback-controlled heating pad (ATC2000, World Precision Instruments). Eyes were protected from dehydration during the surgery with eye ointment (Oculotect Augengel). The scalp overlaying the longitudinal fissure was removed, and a custom head-fixing head-plate was mounted on the skull with cyanoacrylate-based glue (Super Glue Precision, Loctite) and dental cement (Jet Denture Repair Powder) to allow for subsequent head fixation during video-oculographic tracking. Mice were returned to their home cage after anesthesia was reversed with an intraperitoneal injection of flumazenil (0.5 mg/kg body weight; Hameln) and atipamezole (2.5 mg/kg body weight; Antisedan, Orion Pharma) mixture dissolved in saline, and after recovering on a heating pad for one hour.

### Visual stimuli for eye-tracking coordinates

Visual stimulation for eye-tracking coordinates was generated and presented via Python-based custom-made software. The visual stimulus was presented on a “V”-shaped dual-monitor setup positioned 15 centimeters from the eye at an angle of 30 degrees to the mouse. To evoke the optokinetic reflex in *Frmd7* knockout and wild-type mice, we presented a square-wave drifting grating simulating binocular rotation. Drifting gratings were presented in eight trials for 30 seconds at a time with 4 seconds of the gray screen between presentations and were drifted in two different directions along the horizontal axis (0° and 180°; monocular and binocular; parallel and anti-parallel) with a spatial frequency of 0.05 cycles/degree and a speed of 5 degrees/sec.

### Rodent video-oculography

The mouse was placed on a platform with its head fixed to prevent head motion interference (Fig 2A and Fig S2). Head fixation was achieved using a metallic plate implanted cranially. To minimize obstruction of the visual field-of-view, a 45-degree hot mirror was aligned above the camera and lateral to the rodent (Fig 2A_1_). The camera was positioned below the field-of-view (Fig 2A_4_). A monitor for displaying visual stimuli was positioned 15 centimeters from the eye at an angle of 30 degrees (Fig 2A_3_). Behind the monitor, a near-infrared light source was angled at 45 degrees (Fig 2A_2_). A CCD camera (Allied Vision Guppy Pro F-031 ¼” CCD Monochrome Camera) was connected to the PC via a dedicated frame grabber PCIe expansion card (ADLINK FIW62). Using an EyeLoop importer, *vimba*.*py* for Vimba-based cameras, the camera frames were fed to EyeLoop in real-time (∼120 Hz; Fig 1A_1_). Finally, the standard EyeLoop data acquisition module continuously logged the generated tracking data.

**Fig S2:**
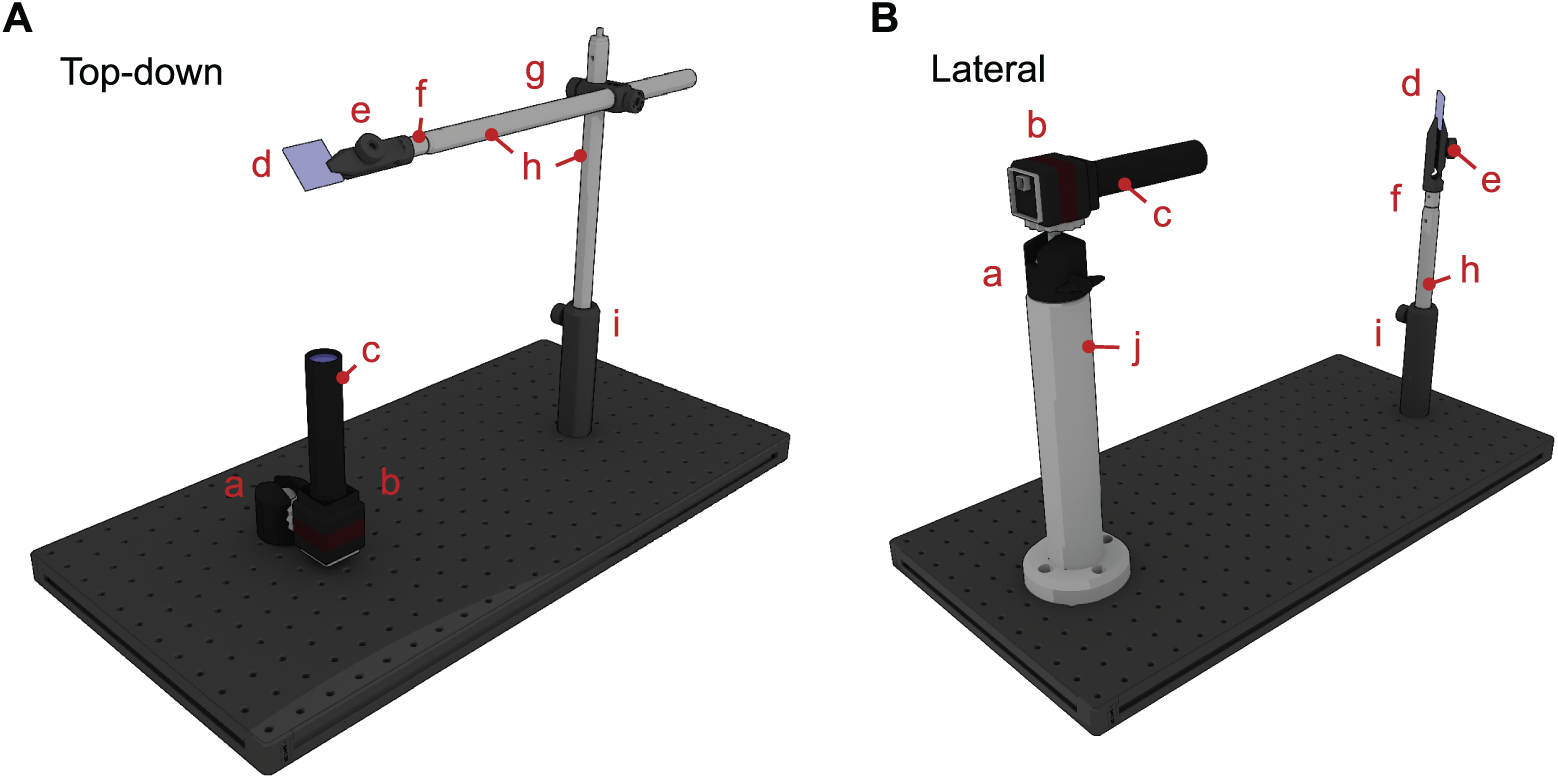
Three-dimensional illustrations of typical video-oculographic setups in research environments. (A) Top-down setting. (B) Lateral setting. a: knuckle-mount (53-887 EdmundOptics). b: camera (Allied Vision Guppy Pro F-031 ¼” CCD Monochrome Camera). c: macro-lens (VZM(tm) 200i Zoom Imaging Lens). d: hot mirror (20mm Square, 45° AOI, Hot Mirror; 62-628 EdmundOptics). e: plate-clamp (PC2 Thorlabs). f: thread adapter (AS4M6M Thorlabs). g: right-angle accessory (RA90 Thorlabs). h: optical posts (TRxV Thorlabs). i: post-holder (PHx Thorlabs). j: damped post (DPxA Thorlabs).

### Human video recording for eye-tracking

The human video sequences were recorded on a human volunteer using a modern smartphone camera set to “slow-motion” mode (240 Hz). Recordings were made in daylight with no infrared or near-infrared illumination source. Illumination was kept static to allow for post hoc angular coordinate calculations based on the corneal reflection position.

### Software

EyeLoop is started via the command line: “python eyeloop.py”. Image data is fed to the system either online (in real-time from a camera) or offline. All parameters may be preset using an “.eyel” configuration file that is passed to the file via argument “-c [path]” or “--config [path]". Alternatively, parameters can be passed as individual arguments (pass “python eyeloop.py –help” for a full list). Note that command-line arguments override “.eyel” configuration presetting. EyeLoop is, by default, tailored to rodent video-oculography (ellipsoid model). To switch to primate mode, pass the argument “-m circular” or “--model circular”. In the following, we describe how to utilize the standard graphical user interface, *minimum-gui*. First, the user selects the corneal reflections by hovering and key-pressing “2”, “3” or “4”. This initiates the tracking algorithm, which is rendered in the preview panel. The user adjusts binarization (W/S) and gaussian (E/D) parameters to improve detection. When the corneal reflections have been satisfactorily selected, the user hovers over the pupil and keypresses “1”. Similar to the corneal reflections, this starts tracking. The user again adjusts binarization (R/F) and gaussian parameters (T/G) for optimal detection. Of note, EyeLoop is capable of filtering out areas of the video sequence crudely by marking them using keypress “b”: Two marks form a rectangular area that is excluded from the tracking algorithm. This feature may be used to filter out obstructing elements. By pressing “v”, marks are undone. For some types of experiments, the angular rotation of the eye is used in real-time. This, however, requires the axes of the video stream and the real-world to be aligned. EyeLoop offers two options for this: Using keys “o” and “p” the video sequence is rotated counterclockwise and clockwise, respectively. In minimum-gui, alignment guidelines are shown in the preview panel. Alternatively, using the angular coordinate module (see Data analyses), the data may be transformed mathematically based on the desired rotation vector. When the tracking settings are deemed optimal, the user initiates the trial by key-pressing “z”, then confirming by “y”.

### Data analyses

#### Quantification of the optokinetic reflex

The optokinetic reflex was quantified by thresholding the first derivative of the eye movements to detect all saccades in the wild-type as described previously [40]. Eye-movement events above the threshold were counted as eye-tracking movements (ETMs). The threshold was adjusted in wild type mice to detect all visually assessed ETMs. Finally, this threshold was applied to the congenital nystagmus disease model mice.

#### Angular coordinates

Because the eye moves rotationally, the two-dimensional video coordinate must be converted to its angular correlate (*r*, θ, φ) as approximated by:

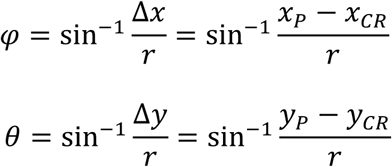

Where *r* is the effective rotation radius. θ and φ are the altitude and azimuth of spherical rotation, respectively. How to calculate the effective rotation radius has been described elsewhere [12]; for reference, mouse *r* is 1.25mm [41], marmoset r is 3.4mm [42, 43], and human r is 6.4mm [44]. EyeLoop includes a standard conversion module that offers rotational matrix transformation to align data to the proper horizontal and vertical axes. Users manually set the conversion module’s animal species via a class argument.

#### Pupillary area

To compute the pupillary area, we must determine the metric unit pupillary radius (Fig S2). As our outset, we calculate the angular coordinates of the pupillary center and the extremum via the angular conversion method described previously (Fig S3.1). By subtraction, we obtain the angular dimension of the radius, Ø. Using the law of sines, we calculate the pupillary radius in millimeters (Fig S3.2):

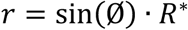

Here, R* represents the effective rotation radius of the eye as described for angular conversion. Finally, we find the pupil area using the basic formula (Fig S3.3):

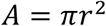

For ellipsoid pupillary models, we perform this algorithm on both dimensions of the ellipse and return the mean. Of note, to scale pupil size calculations, researchers commonly use a reference item placed in the physical plane where the animal is subsequently positioned for eye-tracking [45]. However, this introduces multiple points for error. First, the animal may not be positioned in the exact plane as the reference, making it less valid as a scale. Second, when the eye is not looking straight at the camera, the pupil is slightly distorted due to lens refraction and eye curvature. Our method circumvents these errors using a mathematical model of the eye supplemented by anatomical constants described in the literature. Accordingly, this method may be applied to other animal species than mice, and with no need for calibration.

**Fig S3.**
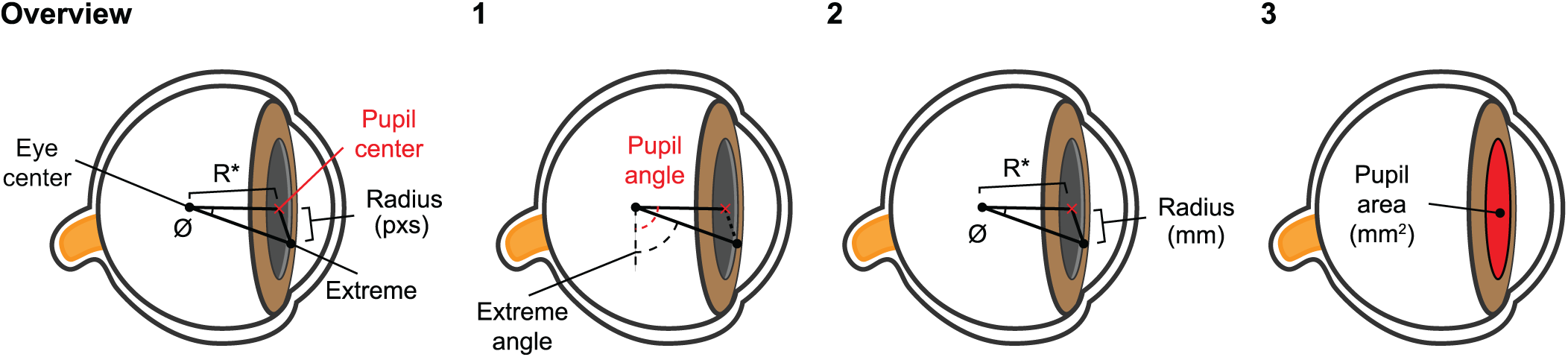
The algorithmic overview of pupillary area calculations.

### Data acquisition

EyeLoop includes a standard data acquisition module. At the end of each frame, all critical data are saved as a timestamped, indexed JSON structure. This data includes the ellipsoid parameters for the pupil and corneal reflection. Additionally, interfacing modules can append to the data output via dictionary calls. Retrieval and parsing of the save log data are possible with common data interchange frameworks, such as JSON. The JSON log may be converted to CSV via EyeLoop’s parser utility. Alignment of the data log to external laboratory components, such as a dedicated visual stimulus or a two-photon microscope, is easily achieved using a dedicated data acquisition device (e.g., National Instruments USB DAQ).

### Parser utility

The parser utility offers an easy method for post hoc analysis of EyeLoop data. This utility includes functions to load the log file, extract timestamps, or unique keys, compute coordinates or pupil area. In addition, it offers converting JSON into CSV.

